# Structural and functional organization of visual responses in the inferior olive of larval zebrafish

**DOI:** 10.1101/2021.11.29.470378

**Authors:** Rita Felix, Daniil A Markov, Sabine L. Renninger, Raquel Tomás, Alexandre Laborde, Megan R Carey, Michael B Orger, Ruben Portugues

**Affiliations:** Champalimaud Research, Champalimaud Centre for the Unknown, Lisbon, Portugal 1400-038; Max Planck Institute of Neurobiology, Sensorimotor Control Research Group, Martinsried, Germany; Institute of Neuroscience, Technical University of Munich, Biedersteiner Strasse 29, 80802 Munich, Germany; Munich Cluster of Systems Neurology (SyNergy), Feodor-Lynen-Strasse 17, 81377 Munich, Germany

## Abstract

The olivo-cerebellar system plays an important role in vertebrate sensorimotor control. According to a classical theory of cerebellar cortex, the inferior olive (IO) provides Purkinje cells with error information which drives motor learning in the cerebellum. Here we investigate the sensory representations in the IO of larval zebrafish and their spatial organization. Using single-cell labeling of genetically identified IO neurons we find that they can be divided into at least two distinct groups based on their spatial location, dendritic morphology, and axonal projection patterns. In the same genetically targeted population, we recorded calcium activity in response to a set of visual stimuli using 2-photon imaging. We found that most IO neurons showed direction selective and binocular responses to visual stimuli and that functional properties were spatially organized within the IO. Light-sheet functional imaging that allowed for simultaneous activity recordings at the soma and axonal level revealed tight coupling between soma location, axonal projections and functional properties of IO neurons. Taken together, our results suggest that anatomically-defined classes of inferior olive neurons correspond to distinct functional types, and that topographic connections between IO and cerebellum contribute to organization of the cerebellum into distinct functional zones.

## Introduction

The olivo-cerebellar system plays a key role in sensorimotor control and coordination in vertebrates. The inferior olive sends climbing fiber projections to cerebellar Purkinje cells, where each climbing fiber makes extensive excitatory synaptic connections with a single Purkinje cell in such a way that a single presynaptic action potential elicits a characteristic Purkinje cell complex spike (Eccles et al., 1966). These complex spikes play a fundamental role in modulating the simple spikes that Purkinje cells fire in response to granule cell input, and thus fine tune cerebellar output (Marr, 1969; Albus, 1971; Ito et al., 1982).

Studies in mammals have identified a population of inferior olive neurons that receives input from direction-selective cells in the accessory optic system, a visual pathway mediating the detection of optic flow (Soodak and Simpson, 1988). This population conveys visual sensory error signals to the cerebellum in the form of retinal slip, giving rise to complex-spikes that are direction-selective, and out of phase with simple-spike firing rate (Ito, 1982; Stone and Lisberger, 1990). In addition to direction selectivity, electrophysiology studies have found evidence of functional organization of climbing fiber projections. For instance, climbing fibers carrying motion signals consistent with rotation about horizontal or vertical axes, project to distinct zones of the cerebellar flocculus (Schonewille et al., 2006; Pakan et al., 2011).

Recent calcium imaging of Purkinje cells in zebrafish larvae has revealed distinct areas of the cerebellum associated with motion stimuli that drive distinct visuo-motor behaviors: the optomotor response (OMR), which drives the fish to swim and turn in the direction of visual motion, and the optokinetic reflex (OKR), which drives the eyes to track the direction of the rotation and make rapid resetting saccades in the opposite direction (Matsui et al., 2014; Knogler et al., 2019). Additionally, electrophysiological recordings from larval zebrafish Purkinje cells, located in different regions of the cerebellum, showed that complex spike responses can be grouped into different visual categories, related with changes in luminance and direction selective translational or rotational motion (Knogler et al., 2019). Since translational and rotational motion stimuli drive different visually-driven motor behaviors in larval zebrafish (Neuhauss et al., 1999), these results have led to the proposed separation of the zebrafish cerebellum into different behavioral modules. According to this model, activity in the medial cerebellum is associated with swimming and turning movements during the OMR, the medial-lateral cerebellum processes changes in luminance, and the lateral cerebellum is involved in eye and body coordination during the OKR (Matsui et al., 2014; Knogler et al., 2019). Distinct functional mapping across medio-lateral cerebellar regions in larval zebrafish is further supported by anatomical mapping of cerebellar outputs (Heap et al., 2013; Takeuchi et al., 2015; Kunst et al., 2019).

Taken together, these studies suggest that spatially segregated and functionally distinct Purkinje cells can differentially modulate activity in distinct brain regions. However, the question still remains as to whether this topographic organization is already present in the inferior olive and fed forward by topography-preserving climbing fiber projections, or whether a different principle organizes the representations in the inferior olive.

Due to technical challenges associated with recording activity from inferior olive neurons in behaving mammals, most functional studies have focused on Purkinje cell complex spikes as a proxy for inferior olive activity. In this study, we take advantage of the larval zebrafish’s small, transparent brain in combination with genetic tools to investigate the inferior olive, and better understand its structure and functional organization at a cellular and population level in response to visual stimuli that drive distinct behaviors. We show that inferior olive neurons can be divided into at least two distinct anatomical types based on their spatial location, dendritic morphology and axonal projection patterns. Furthermore, most inferior olive neurons are direction selective, with functional properties spatially organized within the inferior olive. We describe both anatomical and functional segregation of inferior olive neurons that can be associated with different cerebellar modules, relevant for visually driven behaviors such as the OMR and OKR.

## Results

### Inferior olive cells can be divided into distinct morphological types

To label IO neurons we used the transgenic hspGFFDMC28C GAL4 driver line (Takeuchi et al., 2015), whose pattern of expression within the zebrafish olivo-cerebellar system is limited to inferior olive cells and their climbing fibers, as shown by previous immunohistochemical and anatomical characterization (Takeuchi et al., 2015). By reconstructing morphologies of single IO cells using electroporation or sparse genetic labeling, we confirmed that all IO cells project contralaterally (Figure 1A). We found at least two types of IO neurons can be distinguished based on their dendritic morphology: neurons here referred to as type 1 have bi- or tripolar dendrites arborizing on the medial and lateral sides of the IO; and type 2 neurons with monopolar dendritic trees oriented towards the midline of the brain (Figure 1B). Type 1 and type 2 neurons are similarly represented in the IO (Figure 1D, 19 and 16 cells out of total 53, respectively). Based on registration to a reference brain, we found that these two types of neurons are located in different regions of the IO and have different projection patterns in the cerebellum: type 1 neurons, are mainly located in the caudal-dorsal area of the IO and project to the ventral-lateral area of the cerebellum, whereas type 2 neurons are located in the ventral-rostral part of the IO and project to the dorsal-medial cerebellum (Figure 1C). About one third of labeled neurons (N = 18) had dendritic morphology that could not clearly be placed in one of the two categories and their location and projection patterns lacked any consistent structure, suggesting that they represent different cell types in the IO that are less commonly labeled, or cells that were immature or damaged by the electroporation. These findings show that at least two distinct types of IO neurons can be distinguished based on their dendritic morphology, projection patterns and location within the IO.

**Figure 1.**
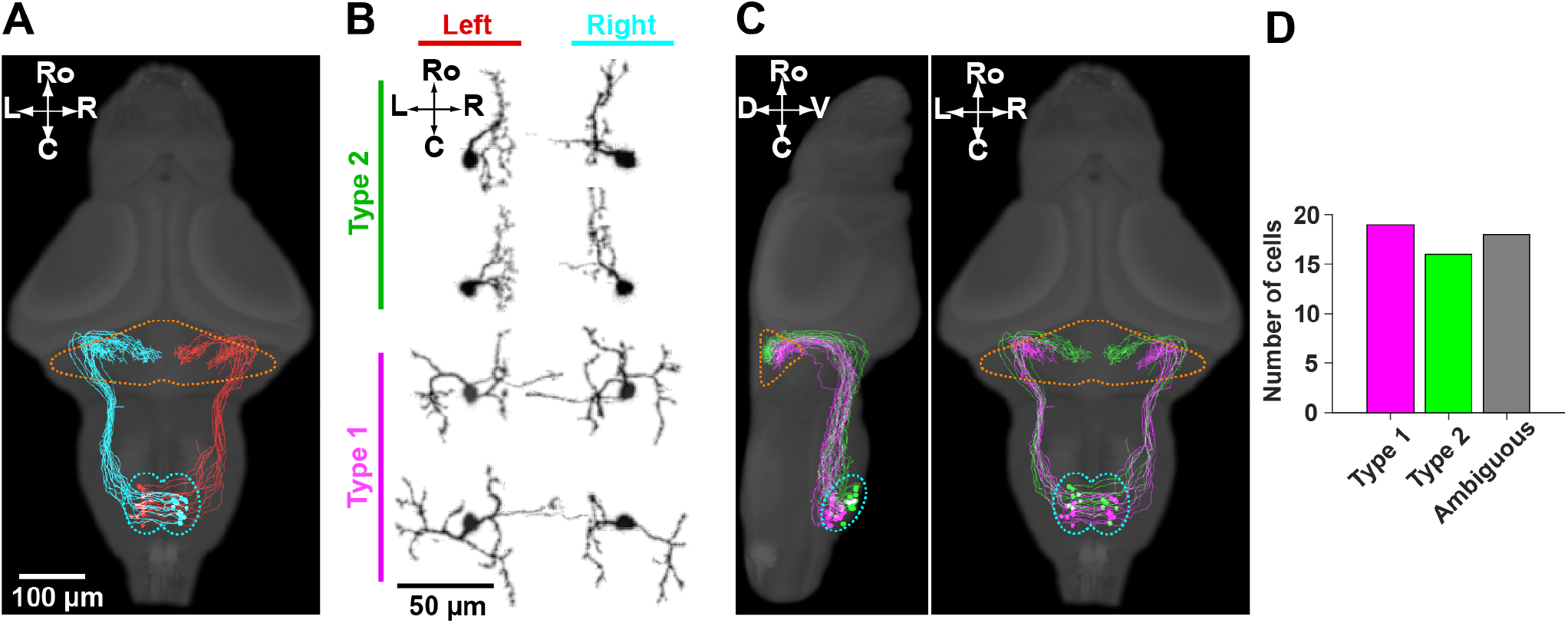
Morphological and anatomical organization of the IO. **IO cells can be divided based on their dendritic morphology, location and projection pattern.** Single cell labelling of inferior olive neurons. **A**. Top view of the larval zebrafish brain showing that inferior olive neurons project contralaterally. Cells located on the right side of the IO (labeled in cyan) project to the left cerebellum and the opposite is true for IO left neurons (labeled in red). **B**. Both left and right inferior olive neurons can be divided in two types based on their dendritic morphology. Type 1 neurons, labeled in magenta, have bi- or tri-polar dendritic trees, while type 2 neurons, labeled in green, are unipolar. **C**. Side and top views of the larval zebrafish brain depicting type 1 and 2 neurons. Type 1 and 2 neurons are located in distinct areas of the IO and project to different regions of the cerebellum. Type 1 neurons (in magenta) are located in the caudal, more dorsal, region of the IO and project to the lateral cerebellum. Type 2 cells (in green) have a more rostral and ventral position within the IO and project to the medial cerebellum. **D**. Quantification of the different neuron types shows that they are similarly represented in our dataset.

### Inferior olive cells are mostly direction selective and are spatially organized

We used the same GAL4 line to drive expression of a GCaMP calcium indicator (GCaMP6fEF05), which incorporates previously described mutations into the GCaMP6f variant (Chen et al., 2013) (Sun et al., 2013), and which we found to give high signal to noise responses in zebrafish neurons. Using 2-photon imaging, we recorded inferior olive activity in response to translational motion, consisting of drifting gratings in 8 different directions, and rotational motion of a radial ‘windmill’ pattern (Figure 2A). These stimuli elicit OMR and OKR, respectively (Supplementary Figure 1A,B). Stimuli were presented in both halves of the visual field on a screen below the fish. Each stimulus presentation took 21.4 s, composed of a stationary period of 6s, followed by a moving period of 10 s (10 mm/s for translational gratings and 22.5 °/s rotational windmill), and stationary again for 5.4 s (Supplementary Figure 1F). For each imaging plane, we presented translational motion of gratings in 8 different directions in a randomized order, followed by clockwise (CW) and counter-clockwise (CCW) rotational motion (Supplementary Figure 1F). Larval zebrafish at 6-7 days post fertilization (dpf) were embedded in low melting temperature agarose, which was then partially removed to allow for tail and eye movements that were recorded with a high speed camera at 700 frames per second (fps) and 100 fps, respectively (Supplementary Figure 1C). We performed manual ROI selection assisted by a custom-written Matlab tool that allowed for 3D identification and quantification of inferior olive cell somata across fish (Supplementary Figure 1D; n = 12 fish, N = 1106 cells). Afterwards, inferior olive cells activity traces were extracted and aligned with visual stimuli and behavior traces (Supplementary Figure 1E).

**Figure 2.**
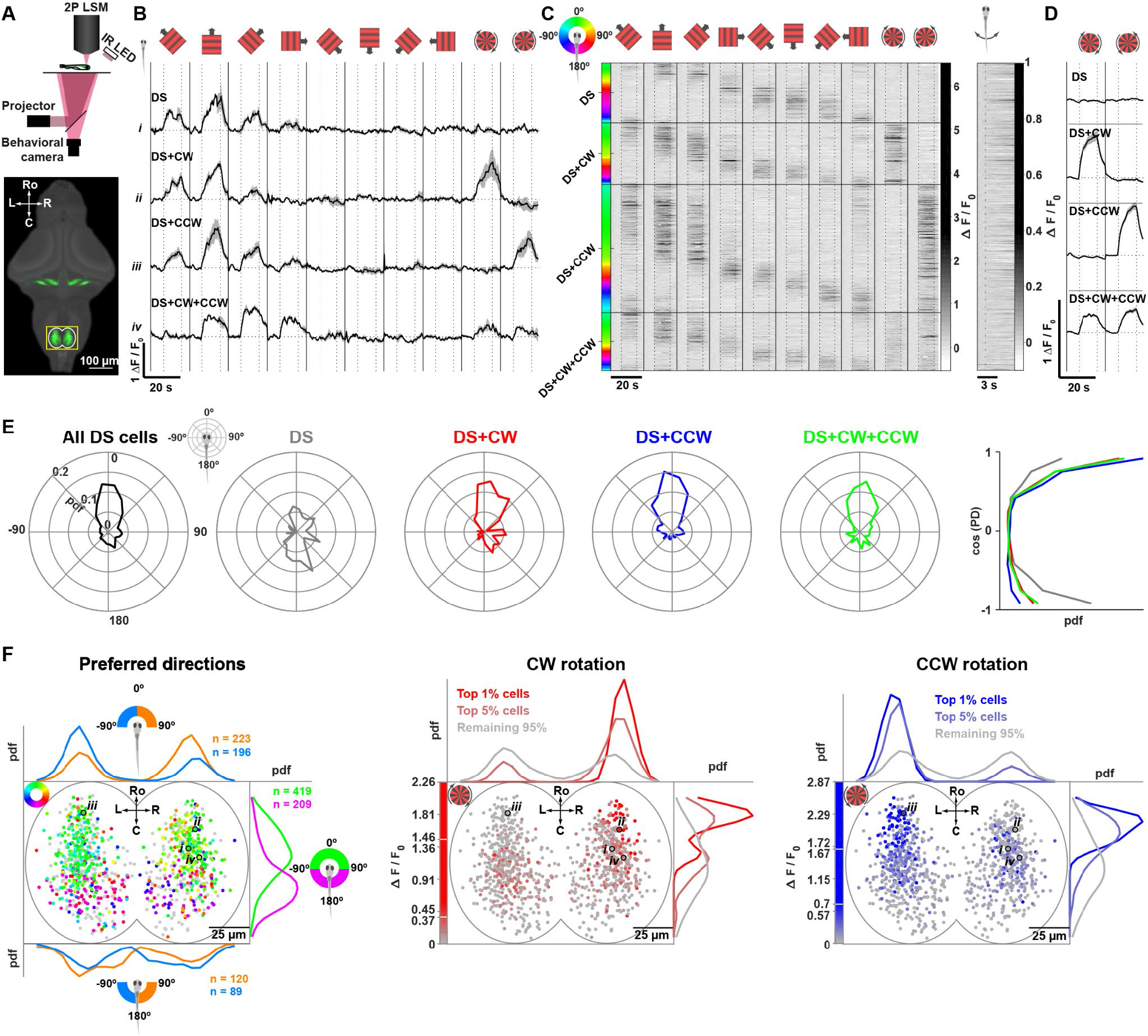
Organization of direction-selectivity in the IO. **A**. Top, schematic of 2-photon microscope used for calcium imaging. Bottom, average larval zebrafish brain reference expressing GFP specifically in IO neurons (hspGFFDMC28C line from Takeuchi et al., 2015). Yellow inset depicts the IO and represents the area imaged during the experiments described below. **B**. Average dF/F responses of forward-selective representative example cells to whole-field translational motion in eight directions and two whole-field rotations (stimulus angle in relation to the fish body is represented on the top row). Cells are described as direction selective (DS) only if they were active exclusively to the translational motion stimuli, or DS+CW or DS+CCW if they also responded to clockwise or counter-clockwise rotation, respectively. DS+CW+CCW represents a population of cells that respond both to whole-field translational and rotational motion (in both directions). Shadows represent SEM. Vertical dashed lines separate stationary and moving periods of the stimulus: stationary (6 seconds), moving (10 seconds) and stationary (5.4 seconds). Horizontal dashed lines represent each cell baseline. Vertical lines separate different visual stimuli. **C**. Left, raster plot for all active cells. Each row represents a cell’s average dF/F response to the ten stimuli shown on top, ordered by preferred direction and separated by a horizontal line for each of the 4 DS categories. On the left, each cell is color-coded for its preferred direction (PD) following the color wheel at the top (green represents forward preferred direction). Right, raster plot for dF/F bout-triggered averages for the same neurons. Dashed line marks bout onset. n = 12 fish, N = 628 cells. **D**. Average dF/F response of all cells in each DS group to clockwise and counter-clockwise whole-field rotational motion. **E**. Distribution of preferred directions (tuning curves) of all cells and of the 4 different groups of cells independently. 0° represents forward, 90° rightward, 180° backward and -90° leftward directions. Probability density function (pdf) as function of cosine of preferred direction for each of the 4 groups. **F**. Left, spatial distribution of preferred directions of inferior olive cells. Each dot represents a cell in a reference inferior olive color-coded for preferred direction as represented in the color wheel, or grey for active cells that were not direction selective. Green and magenta curves show the rostral-caudal distribution of forward and backward preferring cells respectively, separated according to the sign of the PD cosine. Blue and orange curves show the distribution of left and right preferring neurons respectively, separated into a forward preferring (top) and backward preferring (bottom) group. Middle and right, spatial distribution of rotation selectivity of inferior olive cells. Each dot represents a cell in a reference inferior olive, color-coded for dF/F response for CW and CCW rotations in red and blue, respectively. White lines on the color bars denote 99th and 95th percentiles, used for presenting the spatial distributions of top 1% (10 cells), top 5% (48 cells) and remaining 95% (919 cells) active cells. Pdf, probability density function. n = 12 fish, N = 1106 cells, from which 967 were active and 628 were direction selective.

The majority of the identified IO neurons were active in response to at least one stimulus during the experiment (891/1106 cells, 81%). Out of these active neurons, 609 (68%) cells were direction-selective, meaning, these neurons showed larger calcium transients in response to a particular direction of stimulus motion (see Material & Methods). Based on responses to rotational stimuli, we divided all direction-selective neurons into 4 classes. Cells could be described as direction selective only (DS) if they were active exclusively to the translational motion stimuli, or DS+CW or DS+CCW if they also responded to clockwise or counter-clockwise rotation, respectively. Some direction selective cells responded to rotational motion in a non-selective manner DS+CW+CCW (Figure 2B).

Inferior olive DS cells covered the full range of directions we showed, with a higher representation of forward selective cells (Figure 2C, left; 2E, All DS cells). This direction selective activity was locked to motion onset in the preferred direction and didn’t correlate with swim events, as we did not see this kind of data structure when we aligned cells’ responses to bout onset (Figure 2C right). In addition, we did not find orientation-selective cells that responded to motion onset in opposing directions.

Although we could find cells tuned to all directions within each class (Figure 2C), the distribution of preferred directions was different between classes (Figure 2E). Direction-selective cells have an overall preference for forward over backward motion (Figure 2E, All DS cells). Cells that only responded to translational motion (Figure 2E, DS) had a more even distribution of forward and backward preferred directions. On the other hand, direction selective cells that were active during rotational stimuli (CW, CCW or both), mostly showed a forward preferred direction (Figure 2E, DS+CW, DS+CCW and DS+CW+CCW). This is also clear if we look at the distribution as a function of the cosine of the preferred direction, with cosine close to 1 corresponding to forward and -1 to backward preference (Figure 2E, right). Taken together, these results show that the majority of inferior olive cells are driven by visual motion stimuli and are direction selective. The distribution of preferred directions has a large peak around forward motion, and a smaller peak for backward directed motion.

To compare the spatial distribution of the IO cell types, we registered all anatomical stacks of individual fish to a common anatomical inferior olive reference (similar to the yellow boxed inset in Figure 2A) using the Computational Morphometry Toolkit (CMTK) (Rohlfing and Maurer, 2003) (see Materials & Methods for details). DS cells were spatially organized within the inferior olive (Figure 2F). Namely, neurons tuned to forward motion were more rostral (Figure 2F, green distribution), while backward selective cells had a more caudal position within the IO (Figure 2F, magenta distribution). These cells also had a different left-right tuning bias depending on which side of the IO they were positioned. Forward-selective cells showed an ipsilateral bias in their preferred direction, with more cells in the left IO having a leftward bias (Figure 2F, blue distribution), and a rightward bias found in the right IO (Figure 2F, orange distribution). Interestingly, the more caudal backward-selective cells had the opposite trend, showing a bias towards the contralateral direction. Rotational-responsive cells showed a very strong lateralization, with CW rotation responses found predominantly in the rostral right IO, and CCW-responding neurons concentrated in the rostral left side (Figure 2F). These results suggest that direction-selective inferior olive neurons are spatially organized, with rostral cells being more forward-and rotation-selective, and neurons in the caudal region more tuned to backward directions.

### Monocular and binocular contributions to responses of IO neurons

In the larval zebrafish, all retinal ganglion cell axons project contralaterally (Burrill and Easter, 1994). In order to understand how the responses described above could be constructed from these monocular inputs, we looked at the integration of visual information in inferior olive cells by presenting both monocular and binocular visual stimuli (Supplementary Figure 1G).

To quantify whether a responsive IO cell receives sensory information from both or from only one eye, we defined a monocular index for each cell. It was calculated as the difference between significant responses to stimuli presented to the right and left eyes (averaged across repetitions and directions), divided by the sum of these responses (see Material & Methods). This index ranges from -1, which indicates that a cell responded only to left eye stimulation (monocular left cell), to +1, indicating a monocular right cell. An index close to 0 indicates similarly strong responses to stimulation of both eyes (i.e. a binocular cell). We found that some cells have a monocular bias, meaning that they responded more to stimuli presented to one of the eyes, left or right, and that these receive information predominantly from the contralateral eye (Figure 3B; for an example of a monocular left neuron see Figure 3A *i*; for an example of monocular right neuron see Figure 3A *ii*). However, most cells were binocular (Figure 3B) and were evenly distributed throughout the IO (Figure 3B; for examples of binocular neurons see Figure 3A *iii-vi*).

**Figure 3.**
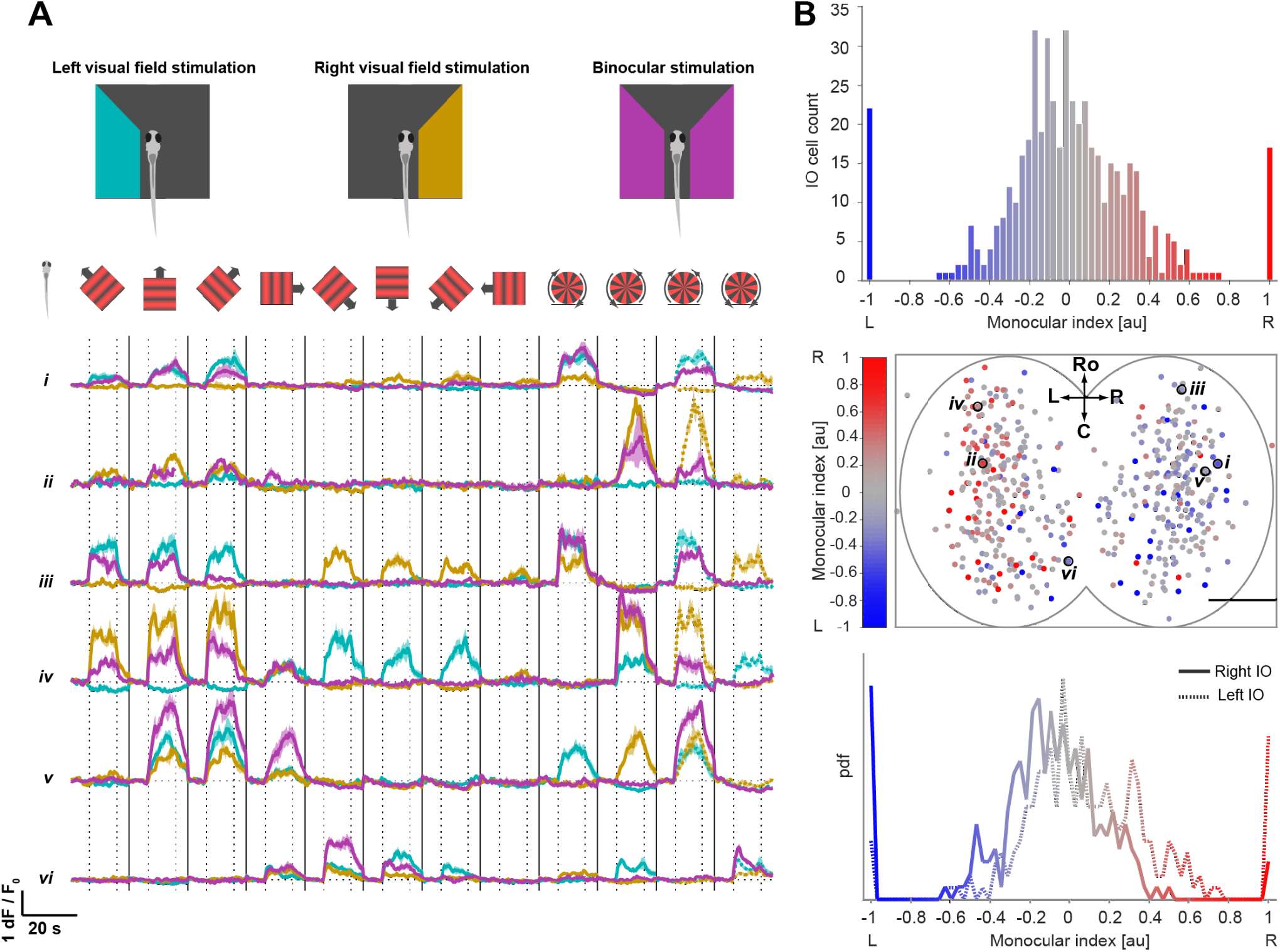
Monocular and binocular responses in inferior olive cells. **A**. Examples of monocular and binocular cells’ responses. Average dF/F responses to monocular stimulation of the left and right visual fields and to binocular stimulation, color coded in cyan, yellow and magenta respectively. Schematic of visual fields stimulation and stimuli angles in relation to the fish are shown on top. Vertical dashed lines separate stationary and moving periods of the stimulus: stationary (6 seconds), moving (10 seconds) and stationary (5.4 seconds). Horizontal dashed lines represent each cell baseline. Vertical lines separate different visual stimuli. *i*. Example cell with monocular left bias. *ii*. Example cell with monocular right bias. *iii,iv*. Example of two cells with opposing translational binocular tuning. *v,vi*. Example of two cells with similar translational binocular tuning. For comparison purposes, all example cells monocular CW and CCW responses are repeated in binocular convergence and divergence stimuli, represented with a dashed line. **B**. Top, distribution of cells’ monocular index. Monocular right-left bias is color coded with a red (right) to blue (left) gradient. Middle, spatial distribution of cells within the inferior olive, color-coded as histogram on top: binocular (unbiased) cells in gray, monocular right bias cells in red and monocular left bias cells in blue. Bottom, probability distribution of monocular index for right (solid line) and left (dashed line) inferior olive cells, showing that inferior olive cells have a monocular bias from the contralateral eye. n = 6 fish, N = 518 cells, from which 511 were active.

We asked which properties of the IO neurons could account for the difference between neurons that responded to rotational motion and neurons that did not. We hypothesized that, in contrast to neurons that did not respond to any rotational motion, neurons that did respond to motion in one sense could have i) different strengths of inputs from the left and right eyes, and/or ii) different preferred directions within left and right visual fields. To test the first hypothesis, that differences in input strength are responsible for preference for rotation direction, we investigated whether the magnitude of the monocular index for each cell correlated with its rotational preference. Although some of the neurons with high monocular index did respond to rotational stimuli, we did not observe a correlation between the magnitude of the monocular index and the response to rotational stimuli presented binocularly (Figure 4A, left). Therefore, differences in monocular bias cannot by themselves explain the variation in rotational selectivity. Binocular rotational motion provides opposing directions of motion to each eye. To test whether responses to rotational stimuli could be explained by different preferred directions within the left and right visual fields, we calculated preferred directions (PDs) of binocular IO cells while only the left or right visual field of the larvae was stimulated (left PD and right PD). We observed that the majority of active binocular IO cells had similar left and right PDs (Figure 4B bottom left and upper right quadrants, and example cells in Figure 3A *v,vi*), and their binocular preferred directions were similar to the monocular preferred directions (i.e. they are green and magenta, respectively). However, we found that a substantial fraction of IO neurons, including those with the strongest responses to rotational stimuli presented binocularly, had opposing PDs between left and right visual fields (Figure 4A right, Figure 4B bottom right and upper left quadrants and example cells in Figure 3A *iii,iv*). These correspond to CW rotation-selective cells and CCW rotational motion selective neurons (Figure 4C).

**Figure 4.**
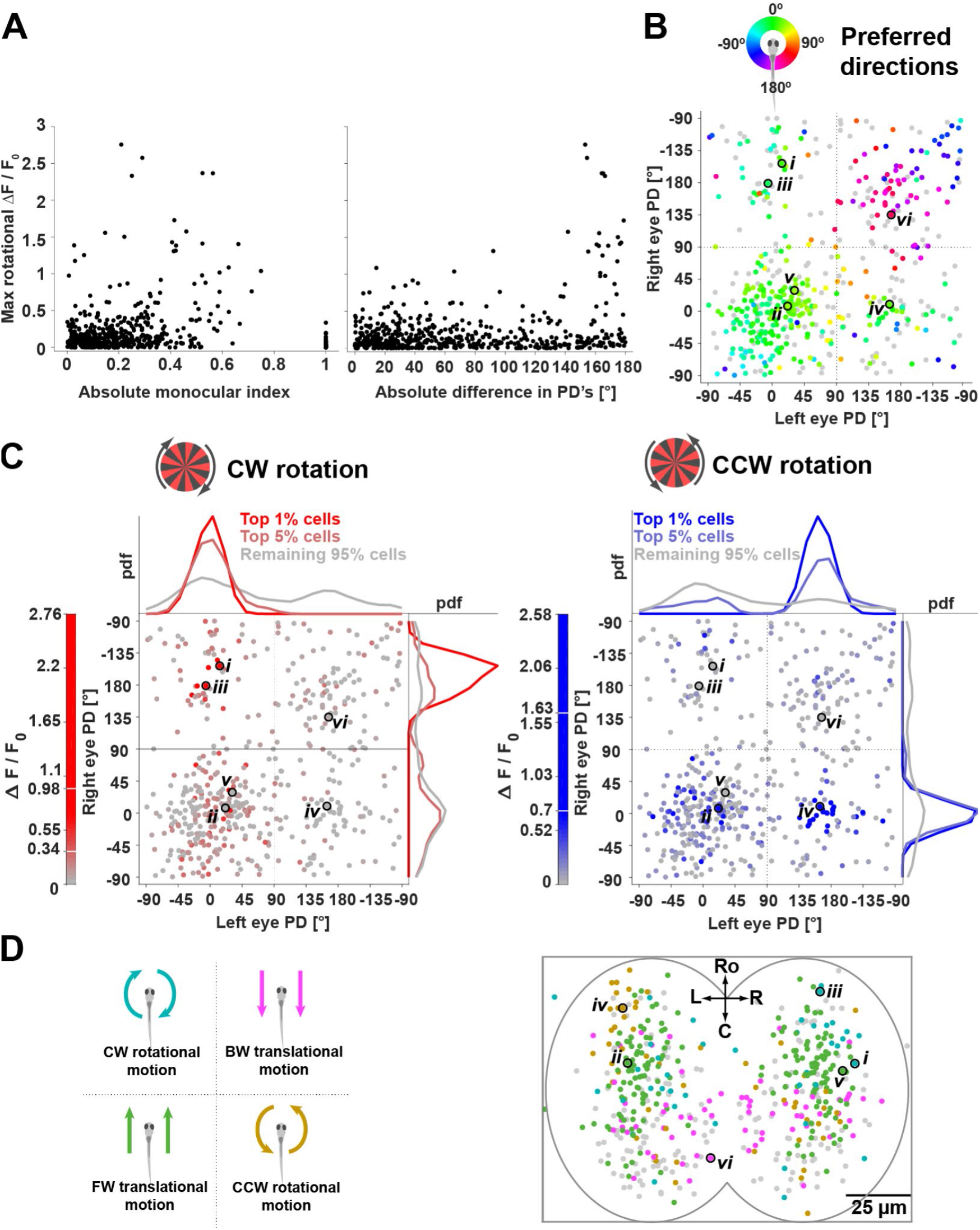
Direction and rotational selective binocular cells are spatially organized. **A**. Left, maximal rotational dF/F responses as a function of the absolute monocular index of each cell (a monocular index close to 1 indicates a monocular cell). Right, maximal rotational dF/F responses as a function of absolute difference in preferred direction of the left and right visual fields. **B**. Binocular cells (binocular) preferred direction as a function of left and right eye preferred direction, color-coded as in color wheel on top. **C**. Left, binocular cells clockwise rotational dF/F responses as a function of left and right preferred directions. Right, binocular cells counter-clockwise rotational dF/F responses as a function of left and right preferred directions. **D**. Binocular cells’ response type as a function of left and right eye preferred directions and their spatial distribution. Translational forward and backward responses are labeled in green and magenta, respectively; responses to clockwise and counter-clockwise rotational motion are labeled in yellow and cyan, respectively. Direction and rotational selective binocular cells are spatially organized within the inferior olive.

The majority of these rotation selective neurons had a forward binocular preferred direction (Figure 4B), which is consistent with the directional tuning properties of DS+CW and DS+CCW within the binocular experiment (Figure 2E). These neurons have opposing preferred directions between the two eyes, but mostly only show responses to forward motion when stimulated binocularly. This suggests the presence of inhibition driven by backward motion on the forward preferring side that leads to binocular rotation responses that are not the simple sum of monocular responses (see example cells in Figure 3A *iii,iv*).

Looking at the distribution of directional and rotational selective binocular cells, we find that they are spatially organized within the IO (Figure 4D), with a pattern consistent with the spatial organization observed before (Figure 2F). Cells with a forward direction selectivity are found more rostrally (green distribution in Figure 2F and Figure 4D), while backward selective neurons had a more caudal location (magenta distribution in Figure 2F and Figure 4D). Rotational selective neurons occupy a rostral and lateral position, with CW rotational selective cells concentrated in the rostral right IO (Figure 2F and cyan population in 4C), while CCW rotational selective neurons are more predominant in the rostral left IO (Figure 2F and yellow population in Figure 4D).

In summary, most inferior olive neurons receive binocular input. A fraction of inferior olive neurons was specifically tuned to rotational motion and had opposing preferred directions between the eyes, i.e. cells strongly responded to CW or CCW rotation if their preferred forward motion was presented to the left or to the right eye, respectively. When larvae were stimulated with translational motion binocularly, these neurons typically preferred forward motion and did not respond to translational motion to the back, even though either the left or right visual receptive field of these neurons was tuned to backwards motion. This suggests that binocular cells do not simply sum their monocular inputs, but that they integrate these non-linearly, in order to compute a behaviorally relevant stimulus feature. Finally, binocular cells were spatially organized within the inferior olive according to their function.

### Different inferior olive functional groups have distinct projection fields in the cerebellum

From our single neuron labeling, we found that two regions could be identified within the inferior olive, referred to as caudal and rostral, that contain neurons with clearly different dendritic morphologies and projection patterns. We were interested in whether this separation of neuron types mapped onto any of the observed functional properties. Therefore, we performed volumetric light-sheet imaging to simultaneously record calcium activity from IO cells’ soma and axons (Figure 5A). Larval zebrafish were paralyzed to avoid potential movement artifacts (Figure 5A) and presented with whole-field translational and rotational stimuli (Figure 5A).

**Figure 5.**
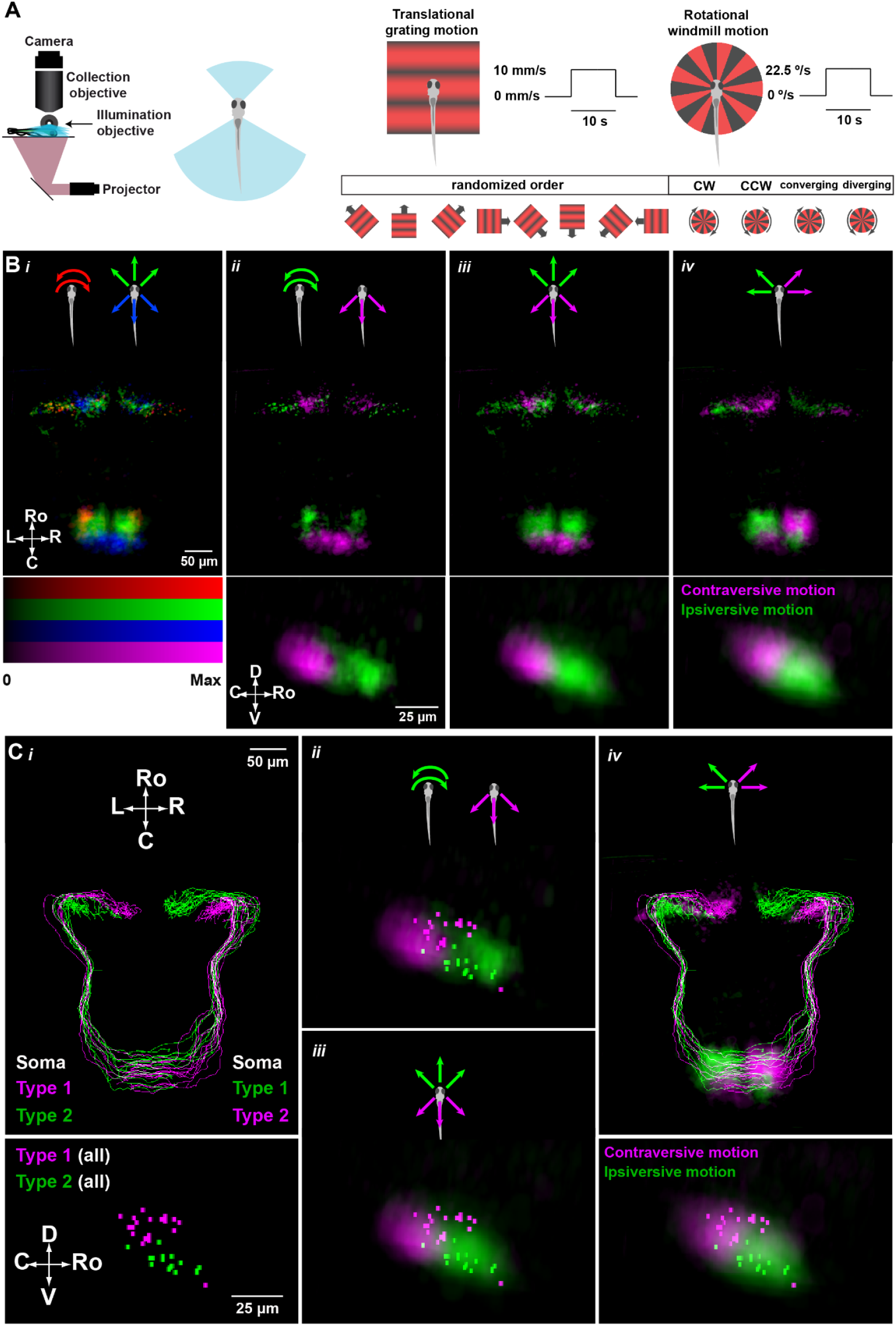
Different anatomy inferior olive groups correspond to different functional types. **Simultaneous imaging of IO and climbing fibre projections. A**. Schematic of light-sheet microscope used for fast volume calcium imaging. Schematic of head-embedded paralyzed larval zebrafish for light-sheet imaging. Whole-field translational sine grating and rotational square windmill visual stimuli that were presented to the fish. Schematic of experimental protocol. The stimulus was presented from below, centered under the fish’s head and each stimulus was presented for 21 s (6 s stationary, 10 s moving, 5 s stationary). Translational gratings moved at 10mm/s and rotational windmill at 22.5 °/s. For each brain volume, we presented 5 repetitions of translational gratings in 8 different directions in a randomized order, followed by clockwise, counter-clockwise, converging and diverging rotational motion. **B**. Light-sheet voxel maps maximal projection of active inferior olive cells’ soma and axons (top) and side view of the inferior olive (bottom) of an average of 28 fish. **B*i***. Projection of the distribution of voxels categorized as forward selective (green), backward selective (blue) or rotation selective (red), averaged across fish. **B*ii***. Distribution of voxels categorized as rotation selective (green), or backward selective (magenta). **B*iii***. Distribution of voxels categorized as forward selective (green), or backward selective (magenta). **B*iv***. *Top*. Distribution of voxels categorized as left selective (green), or right selective (magenta). *Bottom*. Distribution of inferior olive voxels selective for ipsiversive (green) and contraversive (magenta) motion. **C*i***. Single cell projections of type 1 and type 2 neurons (dorsal view; top) and their cell body locations (lateral view; bottom) color coded according to the legends. Note the color code is different between top and bottom panels. **C*ii***. Overlay of type 1 and type 2 neuron soma position and the distribution of inferior olive voxels selective for rotational (green) or backward (magenta) motion. **C*iii***. Overlay of type 1 and type 2 neurons soma position the distribution of inferior olive voxels selective for forward (green) and backward (magenta) motion. **C*iv***. *Top*. Overlay of single cell’s projection maps with map of voxels selective for forward-left/leftwards (green) and forward-right/rightwards (magenta) translational motion stimuli (shown in B*iv*). *Bottom*. Overlay of type 1 and type 2 neurons soma position and the distribution of inferior olive voxels selective for ipsiversive (green) and contraversive (magenta) motion.

As in the previous experiments with binocular visual stimulation, the protocol consisted of translational gratings moving in 8 different directions, in a randomized order, followed by rotational motion in a clockwise, counter-clockwise, converging and diverging manner (Figure 5A). Fish brains were imaged at 100 frames per second, in volumes of 44 planes (2.3 Hz per volume), that covered a 220 µm square area. Active voxels were detected across 28 fish, and registered to the same reference space as the single neuron morphology data. Consistent with the previous two-photon data (Figure 2F), direction-selective activity was observed throughout the inferior olive, and, in each fish, voxels in the cerebellar region were found with similar tunings to those found in the contralateral inferior olive. The pooled data showed the spatial distribution expected from our previous results, with forward-selective DS voxels clustered in the rostral and medial regions of the IO (Figure 5Bi green), and voxels selective for rotation found rostrolateral to these (red). Backward-selective voxels were more caudally located (blue) (see also Supplementary Movies 1, 2 and 3).

The spatial organization of these functional classes within the IO raises the interesting possibility that they could map onto the type 1/2 morphological classes described in Figure 1. Specifically, we would predict that more rostral neurons, responsive to forward and rotational motion, should project to more medial regions of cerebellar cortex, while more caudally-located, backward-preferring neurons would project more laterally. However, analyzing the 3D positions in the IO and cerebellar projection fields of these particular functional groups (Figure 5B ii, iii), we found that the opposite was true: the terminals of rotation selective cells occupied more lateral positions in the cerebellar cortex than terminals of backwards selective neurons (Figure 5B ii). Furthermore, neither of these two types in the olive itself shows a spatial distribution which overlaps well with the morphological classes (Figure 5C ii). Looking at forward vs backward-selective voxels, the forward preferring regions overlap well with the distribution of somata of Type 2 and some Type 1 neurons (Figure 5C iii). However, forward-selective terminals were found with a wide distribution extending both medial and lateral to the backward-preferring domain (Figure 5B iii), also not consistent with a simple correspondence with Type 1 or 2 neurons.

We therefore asked if there was another functional characteristic that more closely matched the Type 1 and 2 morphological pattern. We observed that, within the large population of voxels that responded to motion towards the front half of the fish, there was a spatial organization of their left-right bias, with neurons responding to ipsilateral motion sitting more rostral to those that preferred contralateral motion (Figure 5B iv). Interestingly the terminals of the more rostral domain were located in the cerebellar cortex more medially than those of the caudal group (assuming a crossed projection (Figure 1A)), consistent with the observed Type 1 and 2 anatomies (Figure 5B iv). Superimposing these two functional domains on the arborizations of single neurons (Figure 5C iv top half) and the distribution of somas (Figure 5C iv lower half) showed a good correspondence between these anatomical and functional divisions.

These results suggest the existence of at least 4 spatial divisions within the olivo-cerebellar pathway, solely based on sensitivity to whole-field motion, with characteristic soma distributions and projection patterns. While the organization based on ipsilateral/contralateral bias may correspond to the Type 1/2 anatomical distinction, the rotational and backward motion sensitive groups may be less frequently labeled or missing in the single cell analysis. Taken together, our data shows that anatomically different classes of inferior olive cells are spatially and functionally organized within the inferior olive and in their projection patterns, contributing to the division of cerebellum into different functional zones.

## Discussion

Here we show that inferior olive cells of larval zebrafish can be divided into at least two types based on their dendritic morphology, location and cerebellar projection pattern. Moreover, these anatomically defined classes of inferior olive neurons appear to correspond to distinct functional types, associated with different cerebellar modules relevant for distinct visually-driven behaviors such as the OMR and OKR. Neurons here referred to as type 1 are located in the caudal inferior olive and have bi- or tri-polar dendrites arborizing on the medial and lateral sides of the inferior olive. Type 2 neurons, located in the rostral inferior olive, have monopolar dendritic trees oriented towards the midline of the brain (Figure 1B,C).

In mammals, inferior olive neurons have been shown to be very heterogeneous in their dendritic morphology, ranging between a continuum of ‘curly’ and ‘straight’ morphology types (Vrieler et al., 2019), and are electrically coupled (Llinas et al., 1974; Leznik and Llinás, 2005). According to neurons’ direction oriented dendritic trees and soma localization, it has been suggested that the inferior olive network is organized into areas of densely or weaker coupling but with no clear clustering (Vrieler et al., 2019). One possibility is that bi- or tri-polar inferior olive cells (Type 1) could serve as ‘link cells’ to electrically couple multiple neighboring cells. The dendritic morphology of these Type 1 cells also makes them good candidates for integration of signals from different inputs and possibly convergence of multimodal signals, similar to granule cells (Ishikawa et al., 2015; Knogler et al., 2017).

Beyond dendritic morphology, the two classes of inferior olive neurons we describe exhibited distinct projection patterns, with Type 1 cells projecting to the medial-lateral cerebellum and Type 2 neurons projecting their axons to the medial cerebellar cortex (Figure 1C). Functional imaging and electrophysiology studies have associated the medial and lateral cerebellum regions to the OMR and OKR visually-driven behaviors, respectively (Matsui et al., 2014; Knogler et al., 2019).

We found that the majority of inferior olive cells that we labeled in larval zebrafish were very responsive to visual stimuli (Figure 2B-D). These visual responses were direction-selective and, for the most part, not tightly associated with swimming events (Figure 2C), suggesting a strong sensory component in their activity. Consistent with previous reports describing complex spikes in Purkinje cells, (Knogler et al., 2019; Markov et al., 2021), IO activity was generally phase-locked to motion onset (Figure 2C), suggesting a role of these neurons as ‘sensors’, rather than ‘integrators’, of sensory evidence (Bahl and Engert, 2020; Dragomir et al., 2020; Markov et al., 2021).

Inferior olive cells’ preferred directions covered the full range of directions presented (Figure 2C), but responses tuned to forward or backward directions were relatively overrepresented (Figure 2E). We observed a bias towards forward motion (Figure 2C,E), although previous studies have reported equally well-represented complex spike responses to reverse motion onset (Knogler et al., 2019). This could be due to an approximate twenty-fold increase in the number of recorded cells in our study. It is also possible at this early stage of development that not all IO cell activity that is observed will result in Purkinje cell complex spikes.

The direction-selective visual responses we observed within the inferior olive were spatially organized, with forward and backward preferred neurons located in a rostral and caudal position, respectively (Figure 2F, green and magenta distributions). We also found that inferior olive direction selective neurons had different left-right tuning biases depending on their position along the rostral-caudal and left-right axes. Cells with a forward-left and forward-right bias were located at the rostral left and right inferior olive, respectively; whereas the caudal backward-selective cells followed the opposite left-right trend (Figure 2F, blue and orange distributions). The same was true for rotation selective cells, with clockwise responses more prominent in the rostral right inferior olive and counter-clockwise rotation responses in the rostral left side (Figure 2F).

The functional spatial organization we observed at the inferior olive (Figure 2F) and climbing fiber projection level (Matsui et al., 2014; Portugues et al., 2014; Knogler et al., 2019), suggests topographic organization in connectivity from the inferior olive to the cerebellar cortex. It would be interesting to investigate what are the inputs to the inferior olive and if these are also topographically organized.

Although inferior olive inputs have not been mapped in zebrafish specifically, both mammals and other teleosts have a population of inferior olive neurons that receives input from the accessory optic system or the pretectum, respectively (Brown et al., 1977; Xue et al., 2008; Yáñez et al., 2018). These upstream regions, which receive input from DS retinal ganglion cells, mediate the detection of optic flow. We found that most inferior olive cells were binocular, i.e. they were sensitive to visual information coming from both the left and right eye (Figure 3B). Although binocular rotational selective cells had opposing preferred directions between the eyes, these neurons typically preferred forward motion, and did not respond to backward motion, when stimulated with translational motion binocularly (Figure 4B). Therefore, inferior olive binocular cells’ responses are not simply the linear sum of monocular responses, and inhibition arising from the stimulus to one eye can act to suppress excitatory inputs from the opposite eye. Most pretectal cells are monocular but mechanisms for integration of binocular optic flow have been proposed to be computed at the pretectum circuit level (Kubo et al., 2014; Naumann et al., 2016; Kramer et al., 2019; Wang et al., 2019). Further, binocular integration could happen either within the inferior olive, or in other regions downstream such as the vestibulocerebellum (Masseck and Hoffmann, 2008, 2009).

Taken together, our data shows that a large fraction of inferior olive neurons are sensitive to visual (sensory) motion and encode specific binocular optic flow patterns, resulting in translation- or rotation-selective responses. In addition, we found most inferior olive cells were direction selective and spatially organized according to their function (Figure 2F), suggesting that type 1 and 2 neurons correspond to different functional types. Light-sheet functional imaging in response to the same binocular translational and rotational stimuli allowed mapping of inferior olive responses at the soma and axonal level and show that different functional types have distinct anatomical structure (Figure 5B, C). Our light-sheet functional imaging also shows a population of neurons responsive to whole-field rotational motion that projects to the lateral cerebellum. This population is missing from our single cell anatomical characterization, but is consistent with lateral cerebellar OKR-related activity shown in previous studies (Matsui et al., 2014; Knogler et al., 2019).

Inferior olive cells send topographically specific climbing fiber inputs to Purkinje cells in different cerebellar regions that are associated with different behaviors (turning and forward swimming in the OMR and oculomotor tracking in the OKR). Our functional recordings of the inferior olive neurons that were active during OMR and OKR were extremely consistent across individuals. This stereotypy underscores a remarkable degree of structural and functional organization within the inferior olive and opens up exciting possibilities for further circuit dissection of the olivo-cerebellar system in larval zebrafish.

## Author Contributions

RF and DM conceived and carried out the experiments, and analyzed the data under the supervision of RP, MBO and MRC. SLR and RT developed the GCaMP indicator transgenic line for calcium imaging. AL wrote software for running experiments and processing data. RF, DM, RP, MBO and MRC wrote the paper with input from all authors.

## Acknowledgements

This work was supported by grants to RP from Max Planck Gesellschaft, the Volkswagen Stiftung, the Deutsche Forschungsgemeinschaft (DFG, German Research Foundation) grant PO 2105/2-1 and by the DFG under Germany’s Excellence Strategy within the framework of the Munich Cluster for Systems Neurology (EXC 2145 SyNergy – ID 390857198); to MBO from ERC (NEUROFISH 773012) and the Portuguese Fundação para a Ciência e a Tecnologia (FCT PTDC/NEU-SCC/5221/2014); and to MRC from the ERC (LOCOLEARN 866237) and the Portuguese Fundação para a Ciência e a Tecnologia (FCT-PTDC/MED-NEU/30890/2017). RF was supported by a PhD Fellowship from Fundação para a Ciência e a Tecnologia (SFRH/BD/90105/2012). This work was developed with the support from the research infrastructure Congento, co-financed by Lisboa Regional Operational Programme (Lisboa2020), under the PORTUGAL 2020 Partnership Agreement, through the European Regional Development Fund (ERDF) and Fundação para a Ciência e Tecnologia (Portugal) under the project LISBOA-01-0145-FEDER-022170. Emilio Gualda, Jose Lima, Luigi Petrucco, Lucas Martins and Claudia Feierstein contributed to development of hardware and software for imaging, behavior tracking and data analysis. We thank Herwig Baier, Isaac Bianco, Koichi Kawakami and Miguel Fernandes for sharing constructs and lines.

## Materials & Methods

### Animal husbandry

All experiments were performed on larval zebrafish (*Danio rerio*) at 6-7 days post-fertilization (dpf), unless specified otherwise. The sex of the animals was not determined at this early developmental stage. Experiments were performed in accordance with the European Directive 2010/63/EU and approved by the Champalimaud Ethics Committee and the Portuguese Direcção Geral Veterinária (Ref. No. 019774) and approved protocols set by the Max Planck Society and the Regierung von Oberbayern (TVA 55-2-1-54-2532-82-2016). Zebrafish breeding and maintenance were performed under standard conditions (Westerfield, 2007; Martins et al., 2016). Fertilized embryos were collected in the morning and kept on sets of 20 in E3 medium, at 28° C with a 14/10 h light-dark cycle.

### Transgenic zebrafish lines

For functional imaging experiments, we used the offspring of an incross of a transgenic line expressing a modified version of Gal4 specifically in the IO neurons (hspGFFDMC28C) (rk8Tg; Takeuchi et al., 2015) and a reporter gene UAS:GCaMP6fEF05 (ccu2Tg) for inferior olive specific expression of GCaMP6fEF05. This modified version of GCaMP6f was made by making the mutations D397N/G398A/N399D (Sun et al., 2013) in the CaM domain of GCaMP6f. This version reports activity in zebrafish with better signal to noise ratio (SNR) than GCaMP6f, while maintaining its fast dynamics in comparison to GCaMP6s (Ostrovsky, Renninger et al., in prep.). Imaging experiments used the nacre (mitfa-/-; b692) mutant background, which lack skin melanophores (Lister et al., 1999).

### 2-Photon functional imaging

Functional imaging experiments were conducted using head-restrained preparations of 6-7 dpf zebrafish larvae (Portugues and Engert, 2011). Each larva was embedded in 2% low melting point agarose (Invitrogen, Thermo Fisher Scientific, USA) in a 35mm Petri dish with a Sylgard 184 (Dow Corning) base. After allowing the agarose to set, the dish was filled with E3 medium and the agarose around the tail and eyes was removed to allow for tail and eye movements that were used as a readout of behavior (Supplementary Figure 1C).

The dish was placed onto a light-diffusing screen and imaged on a custom-built two-photon microscope. A Ti-Sapphire laser (Coherent Chameleon) tuned to 950 nm was used for excitation and a custom-written Labview software was used to control the microscope and capture image data. We used a custom-written rendering engine that uses fragment shaders in OpenGL to draw visual stimuli in synchronization with 2-photon imaging software. Visual stimuli were projected from below onto a flat screen at 60Hz using a Optoma ML750e LED projector and a red colored glass long-pass filter (Thorlabs FGL590) and TXRED emission filter (Thorlabs MF630.69) to allow for simultaneous imaging and visual stimulation. Larval brains were systematically imaged from dorsal to ventral, in two micron intervals, at approximately 3Hz (345.6 ms/frame, for a total of 62 frames per stimulus). Each fish was imaged for 30-40 planes which corresponded to approximately 60-80 µm that covered the IO volume.

Two infrared LEDs (850 nm wavelength) were angled between the imaging objective and petri dish to illuminate the fish and allow the tracking of the tail. Behavior was recorded using a Mikrotron EoSens (MC1362) high-speed camera and a National Instruments frame grabber (PCIe-1433). Tail-tracking was performed at 700 frames per second (fps) and eye-tracking at 100 fps using a custom-written software in C#.

### Visual stimuli

For binocular-only experiments we presented the stimuli as shown in Supplementary Figure 1F, namely whole-field sine black and red gratings with a 10 mm spatial period moving in eight directions in a randomized order, followed by whole-field windmill stimulus rotating clockwise and counterclockwise. Experiments that involved both monocular and binocular stimulation had slightly different stimuli as shown in Supplementary Figure 1G. To avoid light contamination of either visual fields during monocular stimulation, the two visual fields were separated by a 1 cm width black patch that was positioned below the fish body (Bianco et al., 2011). In addition, we had a 50° cut off in front of the fish (27.5° in each eye) to prevent stimulation of the fish’s binocular field. To avoid possible light on/off responses in the data recorded, monocular and binocular stimulation were performed in blocks (left, followed by right and finally binocular stimulation) and the first stimulus of the group was presented twice so the first could be discarded. Similar to the binocular-only experiment, each stimulation block consisted of the presentation of translational gratings in 8 different directions in a randomized order, followed by clockwise and counter-clockwise rotational motion; and additional converging and diverging rotational motion for binocular stimulation (Supplementary Figure 1G).

For all experiments, visual stimuli were projected from below onto a flat diffusing screen, and centered under the fish. Each stimulus cycle took 21.4 s (6 s stationary, 10 s moving, 5.4 s stationary), which corresponded to 62 imaging frames (345.6 ms/frame). Translational gratings moved at 10 mm/s and rotational windmill at 22.5 °/s. After each set of stimuli (620 frames/3.6 min for binocular only experiments or 2232 frames/12.9 min for monocular and binocular stimulation), the imaging plane was moved 2 µm ventrally and the set of stimuli was repeated, with translational directions being randomized in each plane.

### Behavior tracking and analysis

In 2-photon functional imaging experiments, tail data was acquired at 700 fps and eye data was acquired at 100 fps. Tail and eye angles were extracted online using custom written software in C#. Briefly, the tail tracking software identified 16 equally spaced points from the middle of the swim bladder to the tip of the tail via a series of radial searches each centered around the previous point. The angles between each pair of tail segments, defined as the lines between two tracked points, were used to compute the cumulative tail angles that were saved to a text file for offline analysis. To track the eyes a point inside each eye was manually identified before the beginning of the experiment. The eye object was then extracted based on the pixel intensity using the watershed algorithm. After that, the orientation of the eye was approximated by calculating the first and second order central image moments of the segmented shape. The eye angle was defined by the angle of the major axis of the approximated eye object relative to the horizontal plane of the image (Hu, 1962). The angles were then corrected to be defined relative to the midline of the fish, and saved to a text file for offline analysis.

Analysis of the behavioral data was performed in MATLAB (MathWorks). Each behavioral trace was interpolated to a new time array with a sampling period of 10 ms, and synchronized with visual stimulus and imaging data. Swim bouts were detected automatically and time stamps for each bout start and end were saved to calculate bout-triggered averages.

### Image analysis

Image analysis was performed with MATLAB (MathWorks). Volumetrically acquired two-photon data frames were first aligned within a plane then across planes to ensure that stacks were aligned to each other with subpixel precision. Any experiments during which the fish drifted significantly in z were stopped and the data discarded. If a frame could not be aligned to adjacent ones due to movement artifacts, it was replaced with NaNs. To generate anatomy stacks for each larva, all aligned frames within each plane were averaged. We then registered the anatomy stacks of individual fish to a common inferior olive reference using the CMTK (Rohlfing and Maurer, 2003).

### Manual cell annotation

Inferior olive cells were selected using a manual 3D ROI selection tool custom-written in MATLAB (MathWorks). This tool loaded an anatomy stack maximum intensity projection and allowed for ROI selection across planes (plane selection example in Supplementary Figure 1D). Given the anatomy of IO cells, we could select with a high degree of certainty a ROI that corresponded to a single cell. The selection was based on an individual level, by clicking on the center of a cell soma, defining its maximum pixel size and automatically selecting similar intensities that could be manually refined to better match the cell’s anatomy. This selection covered the same region in z, so the same cell could be identified and selected in adjacent planes. Since the scanning was performed plane-by-plane, the number of trials for which the activity of a given cell was sampled depended on the number of planes covered by that cell. As a result, different cells might have different numbers of repetitions. The centroid of the middle plane of each cell was used as a proxy for the cell center, and its coordinates were used for inferior olive spatial distribution analysis.

### Data analysis

Inferior olive cells’ activity traces were extracted from manually selected cells. Fluorescence signals from these cells were converted to a (ΔF)/F0 or (dF/F0) value. For each stimulus, dF/F0 was calculated using the fluorescence signal during the time the stimulus was moving (F) and normalized by the baseline, defined by the previous fluorescence signal when the stimulus was stationary (F0). The mean dF/F0 in the window where the stimulus was moving, averaged across trials, was chosen as the response of a neuron to the stimulus. To test for response to motion, we compared cells’ response to a threshold, defined by the mean baseline response plus 2 standard deviations, computed across all baseline frames. Only neurons whose response to at least one stimulus was higher than this threshold were considered active cells. Direction selectivity was quantified by a ratiometric direction selectivity index (DSI), calculated as the vector sum of mean responses to all directions divided by the total response for that cell. This means that a cell that only responded to a stimulus would have a DSI of 1 and an active cell that responded equally to all directions would have a DSI of 0. Preferred direction corresponded to the DSI angle.

Monocular cells were classified as such if they were active during stimulation of one of the visual fields and to binocular stimulation. Cells were classified as binocular if they were active to both left and right visual fields stimulation and/or binocular stimulation. A monocular index was calculated as the absolute difference between responses to stimuli presented to the left and right eyes averaged across repetitions and directions and divided by the sum of these responses, for each cell. To avoid divisions by near-zero values in this computation, we only included responses that were at least two standard deviations greater than the average baseline fluorescence.

Since swimming bouts were recorded at a rate faster than our imaging frame rate, we considered the dF/F of the time point closest to (and after) a bout start, as bout onset to calculate bout-triggered averages.

### Light-sheet functional imaging

Light-sheet functional imaging experiments were conducted using head-restrained preparations of 6-7 dpf zebrafish larvae (Portugues and Engert, 2011). Each larva was paralyzed in bath-applied α - bungarotoxin 2 mg/ml for 5-10 seconds, and embedded in 1.6 % low melting point agarose (Invitrogen, Thermo Fisher Scientific, USA) in a 35mm Petri dish with a Sylgard 184 (Dow Corning) base. After allowing the agarose to set, the dish was filled with E3 medium and the agarose around the eyes was removed to avoid light scattering (Figure 5A). Each petri dish was cut and fit with a cover slip window that was systematically positioned on the left side of the fish. This allowed for unperturbed entry of the focused light sheet laterally onto the fish. The dish was placed onto a light-diffusing screen and imaged on a custom-built light-sheet microscope (Figure 5A). We used a blue excitation laser (MBL-FN-473, 473 nm, 200 mW, Changchun New Industries Optoelectronics Technology), controlled by an acousto-optic modulator (MTS110-A3-VIS, AA optoelectronics) that allowed for rapid changes in light sheet power. The beam passed onto a first 1D galvo mirror (GVS011, Thorlabs) which scanned horizontally in order to create a sheet of light. We used a pair of lenses (AC254-100-A-ML, Thorlabs) to focus the light sheet onto a second 1D galvo mirror that allowed for the scanning of the light sheet vertically through the fish, and illuminate a series of optical slices to perform volumetric imaging. A 1D line diffuser was added in the pathway of the light sheet to reduce horizontal striping in the image due to shadowing of the light sheet by skin structures or blood vessels (Taylor et al., 2018). Fluorescence light was collected with a 20x water immersion objective (XLUMPlanFLN 20x/1.00W) and two 525 nm bandpass filters were used to exclude any non-green light from being detected by the camera (ORCA-flash 4.0, Hamamatsu). Lateral pixel size was measured to be 0.65 µm. Visual stimuli were projected from below onto a flat screen using a laser projector (SHOWWX+ Laser Pico Projector, MicroVision). Fish brains were imaged at 100 frames per second, in volumes of 44 planes (2.3 Hz per volume), that covered an 220 µm axial range. A custom-written GUI developed by José Lima and Lucas Martins was used to control the microscope and capture image data.

For light-sheet binocular experiments we presented the stimuli as shown in Figure 5A. Namely, whole-field sine gratings moving in eight directions in a randomized order, followed by clockwise, counterclockwise, converging and diverging whole-field windmill rotational motion (Figure 5A). Visual stimuli were projected from below onto a flat diffusing screen, and centered under the fish. Each stimulus was presented for 21 seconds (6 s stationary, 10 s moving, 5 s stationary). Translational gratings moved at 10 mm/s and rotational windmill at 22.5 °/s. Each set of stimuli was presented 5 times, with translational directions being randomized in each repetition. Before the experimental protocol started, fish were habituated to laser scanning for 5 minutes. A single green frame, which bled through the filters sufficiently to be detected by the camera, was presented in the beginning and end of the experimental protocol, to allow imaging data and visual stimulus synchronization.

### Analysis of light-sheet data

Image analysis of light-sheet data was performed with MATLAB (MathWorks). Volumetrically acquired light-sheet data was first corrected for rigid translational drift over the course of the experiment identified using the Matlab *imregtform* command. Any experiments during which the fish drifted significantly in z were discarded. We then registered the anatomy of individual fish to reference brain using the CMTK (Rohlfing and Maurer, 2003). For each stimulus, mean response (delta F/F_0_) was calculated by averaging frames during the moving period of the stimulus (F), and normalizing by the baseline (F_0_), which was calculated by averaging the frames when the stimulus was stationary. Each stimulus response was then averaged across the 5 repetitions (trials). This allowed us to create stack average activity maps for each stimulus for all fish. For voxelwise analysis, we selected voxels above a threshold brightness (2 gray values above background on average), which included the inferior olive and its projections. Voxels were placed into response categories using thresholds selected based on the response strengths observed in the data. For forward and backward preference, we considered only the six stimuli with a forward or backward component and defined two categories: 1) Voxels with a forward motion preference and an average response of at least 50% over the 3 forward stimuli. 2) Voxels with a backward motion preference and an average response of at least 25% over the 3 backward stimuli. To compare left-right bias, we considered left, right and forward-left, forward-right stimuli, and selected voxels with a left-right bias and an average response of at least 25% in the preferred direction. Rotation-selective voxels were selected based on a response >75% to one rotational direction and <25% to the other. To display 2D projected maps of the response distributions, we summed the number of voxels in each category. Since the number of voxels in the IO was much larger than the sparser cerebellar projections, the top and bottom halves of each image, which corresponded to climbing fiber projections and inferior olive cells’ soma, respectively, were normalized independently to the maximum observed value.

### Single cell labeling

Labeling of individual inferior olive cells was achieved by using either single-cell electroporation or by outcrossing the Tg(hspGFFDMC28C); Tg(UAS:GFP) (Takeuchi et al., 2015) to a sparse UAS:epNtr-tagRFP reporter line (line mpn123 generated by Miguel Fernandes and Herwig Baier at MPI Neurobiology, Munich). The offspring of such an outcross had very sparse expression of RFP, often in only one or two IO cells, which was ideal for cell tracing.

For single-cell electroporation, 5-6 dpf Tg(hspGFFDMC28C); UAS:mCherry larvae were embedded in 1.5% low melting point agarose, anesthetized in bath-applied solution of MS-222 (tricaine) at a concentration of 0.16 g/L, and inferior olive cells were electroporated under a confocal microscope (LSM 780, Carl Zeiss, Germany) as described previously (Tawk et al., 2009). Briefly, a fine borosilicate glass electrode with filament (final tip diameter ∼ 1 µm) was filled with plasmid DNA pCS2-GAP43-GFP construct (provided by Isaac Bianco) at a concentration of ∼1 µg/µl in distilled water and manipulated through the tissue to a target fluorescent inferior olive cell using a micromanipulator (Sutter Instruments, Novato, CA). 1 - 3 square trains of electric pulses with frequency of 200 Hz, duration of 1s and magnitude of 20-30 V were applied to inject DNA constructs into a single cell using an Axoporator 800A (Molecular Devices, Silicon Valley, CA). After electroporation, dishes with embedded larvae were gently washed with Danieau’s solution (58 mM NaCl, 0.7 mM KCl, 0.4 mM MgSO4, 0.6 mM Ca(NO3)2, 5 mM HEPES buffer) three times to remove the tricaine. After washing, larvae were released from the agarose and allowed to recover in Danieau’s solution overnight.

To image the labelling results, 6 or 7 dpf larvae were anesthetized using tricaine and embedded in 1.5% low melting point agarose. Labelled cells were imaged using a confocal microscope (LSM 780, Carl Zeiss, Germany). For successfully labelled larvae, two stacks were acquired: one for visualizing the cell body and dendritic arbors at higher magnification (Figure 1B), and another to capture the whole span of axonal projections at lower magnification (Figure 1A).

To allow for subsequent anatomical registration (see below), stacks with axonal projections were acquired in two channels. One channel containing dense signal from the majority of the IO cells and their projections (GFP for labelling using sparse expression and mCherry for electroporations) served as an anatomical reference for registration. Another channel contained one or two labelled IO neurons (RFP for labelling using sparse expression and GFP for electroporations). After anatomical registration, axonal projections were traced and reconstructed using the ‘Simple Neurite Tracer’ FIJI plugin (Longair et al., 2011; Schindelin et al., 2012).

### Anatomical registration

To represent functional and anatomical data acquired from different larvae within a common coordinate system, all datasets were registered to one of the two common reference stacks. Anatomical registration was performed using affine volumetric transformation computed by the CMTK (Rohlfing and Maurer, 2003).

The first common reference stack included IO and the cerebellum to cover location of the IO cell bodies and their anatomical projections (IO-CF reference, e.g. Figure 5B). To obtain the IO-CF reference stack, reference channels of the confocal stacks acquired for single cell labelling (see above) were registered to one of these stacks and then averaged. We used the IO-CF reference to register not only the single-cell anatomical data (N = 39 fish) but also all light-sheet functional imaging data (N = 28 fish, Figure 5). Some light-sheet examples could not be registered well to the selected template, and, in these cases, we registered these to the other individual fish that were well registered, and chose the best match. This process was repeated until all functional imaging data was registered to the IO-CF reference.

The second reference stack represented an IO subsection of the IO-CF reference (IO reference, similar to the yellow inset in Figure 2A). It was used for anatomical registration of the 2-photon functional imaging datasets, where CFs were not imaged (N = 18 fish, 12 fish from binocular experiments and 6 fish from monocular and binocular experiments).

Finally, to represent the anatomical organization of the data in the context of the whole brain (e.g. Figure 1A), IO-CF reference was registered to a whole brain reference stack that was previously acquired in the Portugues laboratory by co-registration of 23 confocal z-stacks of zebrafish brains with pan-neuronal expression of GCaMP6f (Tg(elavl3:GCaMP6f; a12200Tg)) (Wolf et al., 2017). To perform this registration, a confocal stack of Tg(elavl3:GCaMP6s (a13203Tg); hspGFFDMC28C (rk8Tg); UAS:mCherry) (Kim et al., 2017) was used as an anatomical link, as it contained both pan-neuronal signal (elavl3) and IO-CF-specific signal (hspGFFDMC28C) in different colors.

## Supplementary figures

**Supplementary Figure 1.**
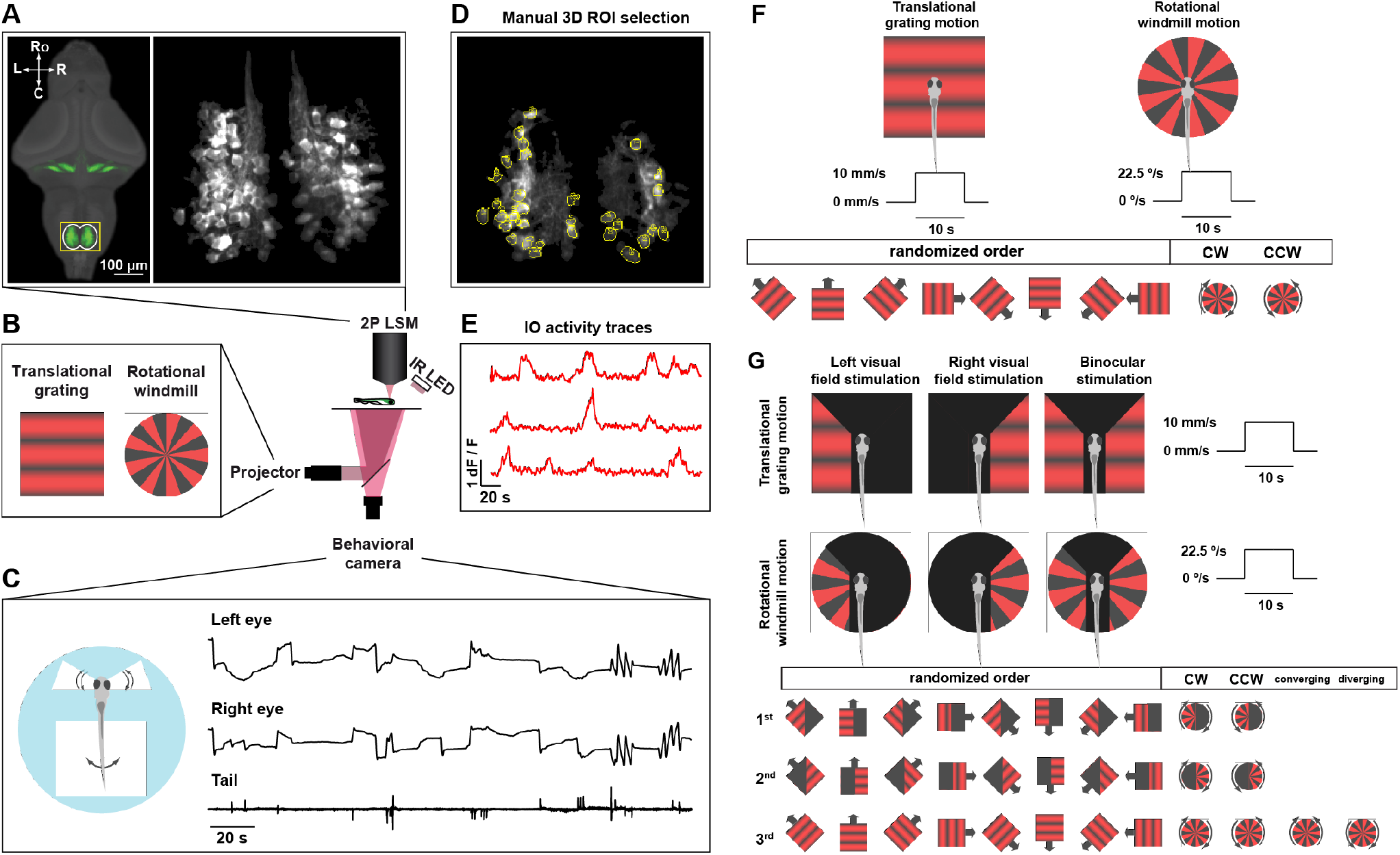
Two-photon setup allows calcium recordings in genetically identified IO neurons with simultaneous tracking of tail and eye movements in response to whole-field translational and rotational motion. Schematic of 2-photon microscope used for calcium imaging (middle). **A**. Left, average larval zebrafish brain reference expressing GFP specifically in IO neurons (hspGFFDMC28C line from (Takeuchi et al., 2015)). Yellow inset depicts the IO and represents the area imaged during the experiments described below. Right, Z-stack maximum projection of a representative fish expressing GCaMP6fEF05. **B**. Overview of the visual stimuli presented to the fish: translational grating and rotational windmill. **C**. Schematic of head-embedded fish with eyes and tail free and an example of recorded behavior traces: left eye, right eye and tail angles. **D**. Example of manual 3D ROI selection in one imaging plane. **E**. Example IO cells activity traces extracted after ROI selection. **F**. Schematic of whole-field translational sine grating and rotational square windmill stimuli that were presented to the fish. The stimulus was presented from below, centered under the fish head and each stimulus cycle took 21.4 s (6 s stationary, 10 s moving, 5.4 s stationary). Translational gratings moved at 10mm/s and rotational windmill at 22.5 °/s. For each imaging plane, we presented translational gratings in 8 different directions in a randomized order, followed by clockwise and counter-clockwise rotational motion. **G**. Schematic of whole-field translational sine grating and rotational square windmill stimuli that were presented to the fish during left, right or left and right visual fields stimulation. To avoid light contamination of either visual field during monocular stimulation, the two visual fields were separated by a vertical 0.5mm black patch that was positioned below the fish body. In addition, we had a 50° cut off in front of the fish (27.5° in each eye) to prevent stimulation of the eyes’ binocular zone. The stimulus was centered under the fish head and each stimulus cycle took 21.4 seconds (6 s stationary, 10 s moving, 5.4 s stationary). Translational gratings moved at 10mm/s and rotational windmill at 22.5 °/s. To avoid possible light on/off effects in the data, monocular and binocular stimulation were performed in groups: left, followed by right and finally binocular stimulation. Translational grating direction was randomized within each block and whole-field rotational stimuli were presented at the end of the 8 directions, in clockwise and counter-clockwise order. Binocular stimulation also included converging and diverging whole-field rotational motion. After each set of stimuli, the imaging plane was moved 2 µm ventrally and the set of stimuli was repeated, with translational directions being randomized in each plane.

